## Supplemental Figures 1-5 for "A Novel Imaging Method (FIM-ID) Reveals that Myofibrillogenesis Plays a Major Role in the Mechanically Induced Growth of Skeletal Muscle"

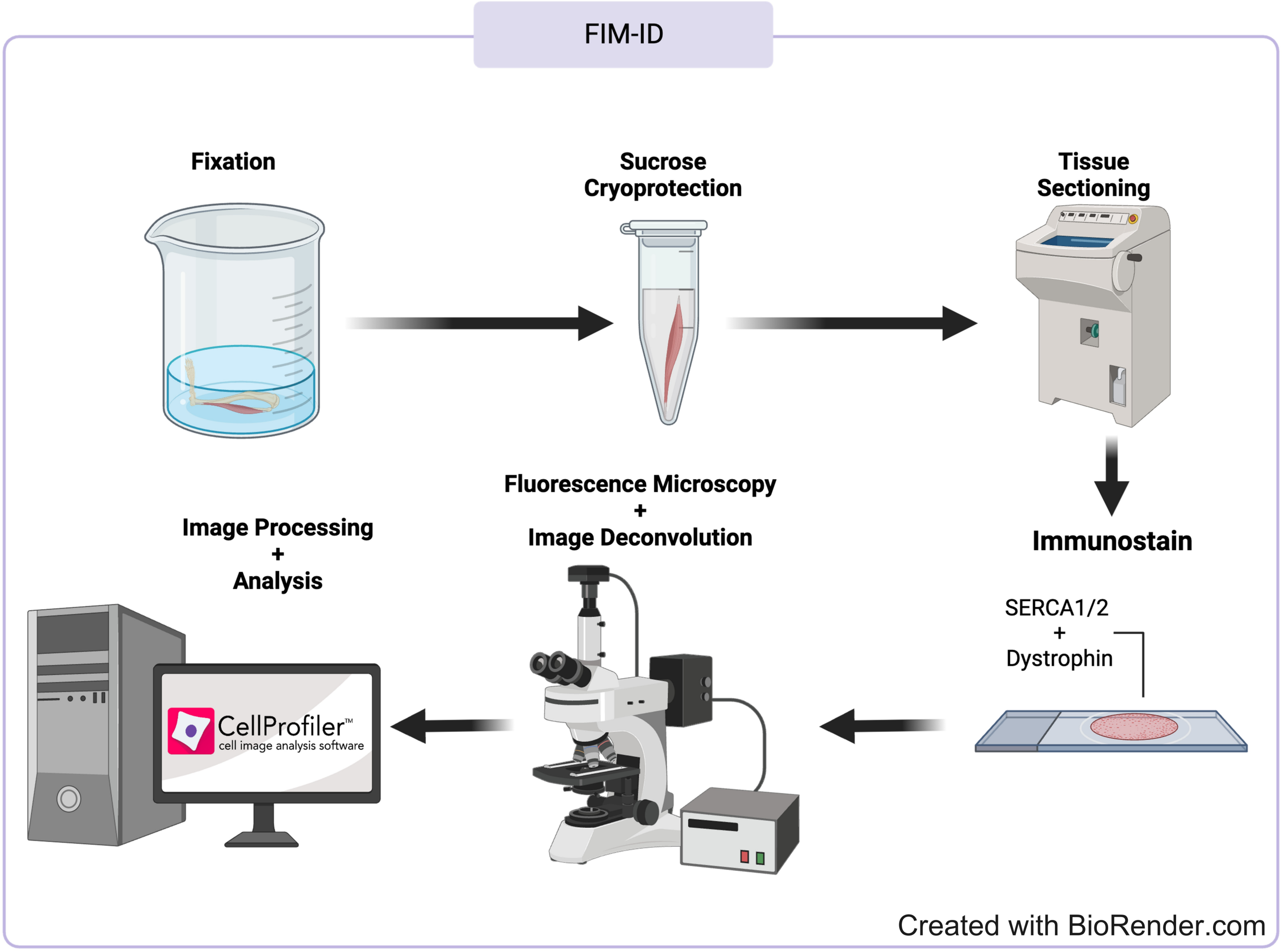


**Supplemental Figure 1. Overview of the FIM-ID Workflow.**

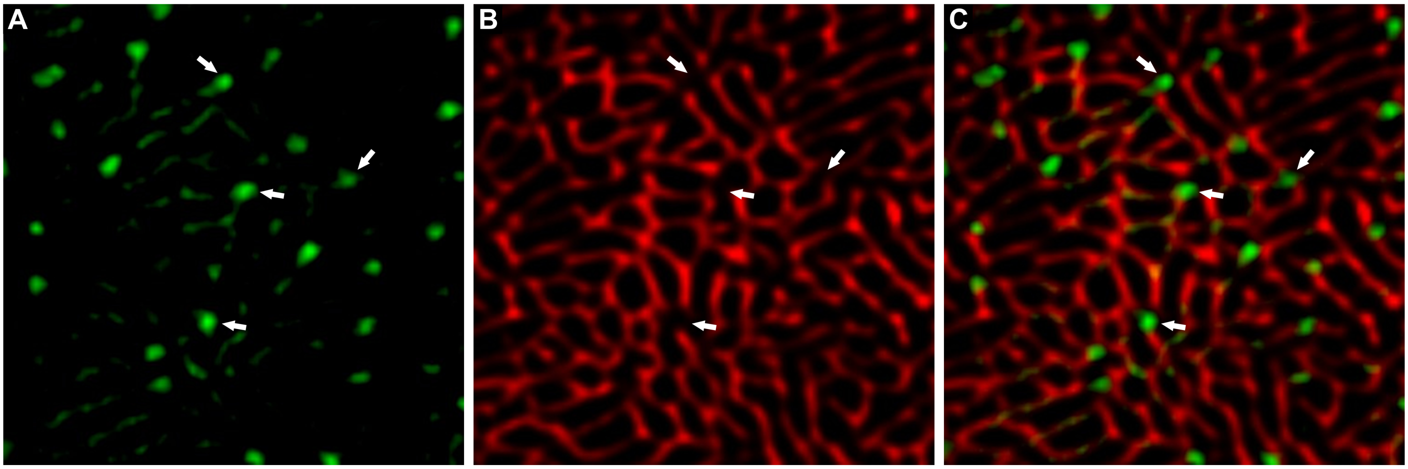


**Supplemental Figure 2. Punctate sites of autofluorescence align with gaps in the signal for SERCA1.** Representative images from a plantaris muscles that was processed for FIM-ID as described in Figure 4. The representative images are from an oxidative (Ox) fiber. The signal for autofluorescence is shown in (**A**) and the signal for SERCA1 is shown in (**B**). The merge of these signals (**C**) reveals that the intense autofluorescent puncta which are frequently observed in Ox fibers are positioned at points in which there is a gap in the signal for SERCA1.

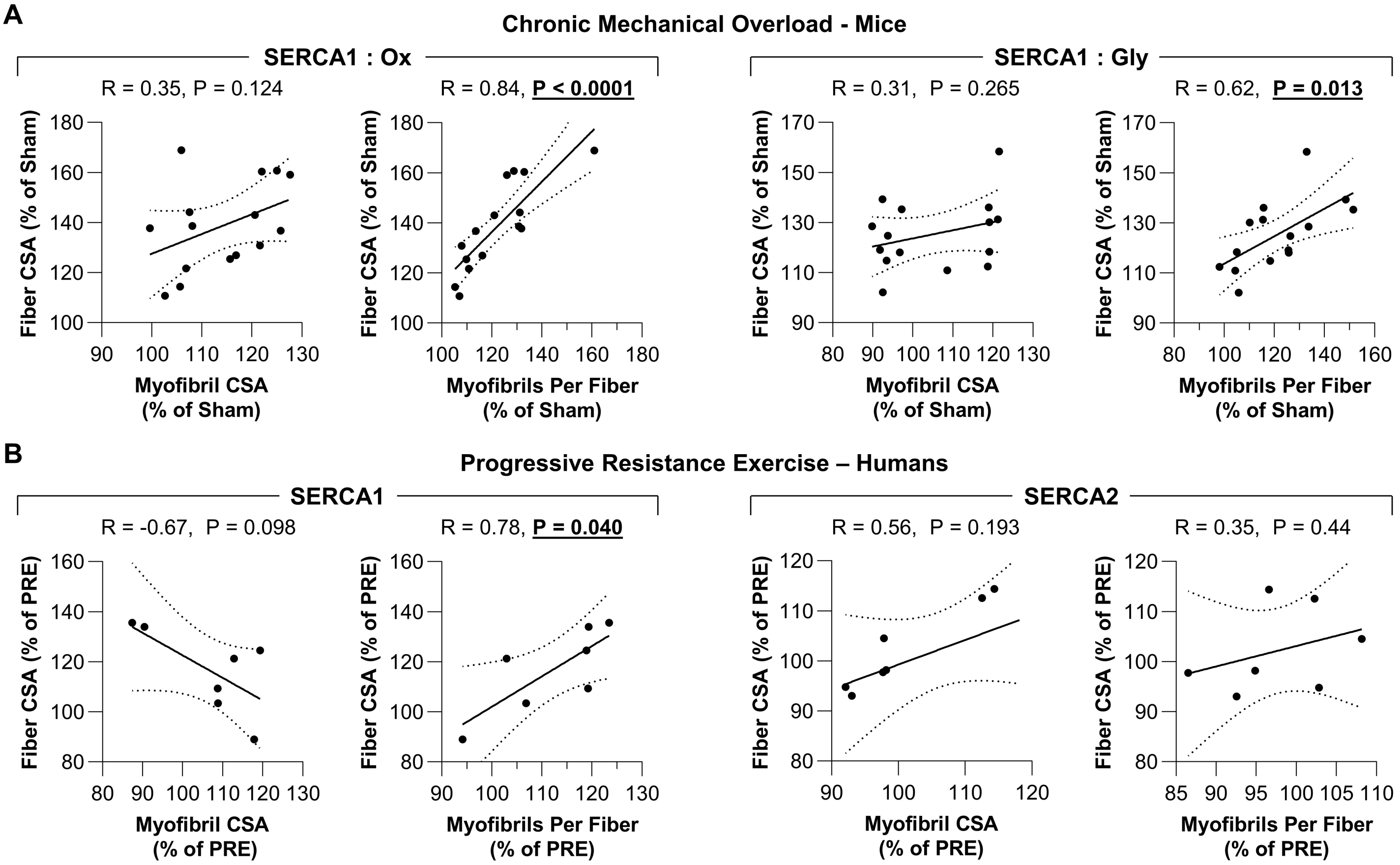


**Supplemental Figure 3. The radial growth of fibers that occurs in response to an increase in mechanical loading is correlated with the induction of myofibrillogenesis.** (**A**) The plantaris muscles of mice were subjected to a chronic mechanical overload (MOV) or sham surgery and then collected after 16 days of recovery. (**B**) Biopsies of the vastus lateralis muscle were collected before (PRE) and after (POST) human participants performed 7 weeks of progressive resistance exercise. Each sample was subjected to FIM-ID and then individual fiber types were analyzed for the mean fiber cross-sectional area (CSA), mean myofibril CSA, and the mean number of myofibril per fiber as reported in Figures 5 and 6. Data for the mouse MOV samples were expressed relative to the mean of the sham samples (% of Sham). Data for each human POST sample was expressed relative to its respective PRE sample (% of PRE). The resulting data were used to create scatter plots that illustrate the relationship between the changes in fiber CSA and myofibril CSA for each sample, or the relationship between the changes in fiber CSA and the number of myofibrils per fiber (i.e., myofibrillogenesis) for each sample. Solid lines represent the best fit from linear regression, the dashed lines represent the 95% confidence intervals, R indicates Pearson's correlation coefficient, and P indicates the likelihood that the relationship is significantly different from zero.

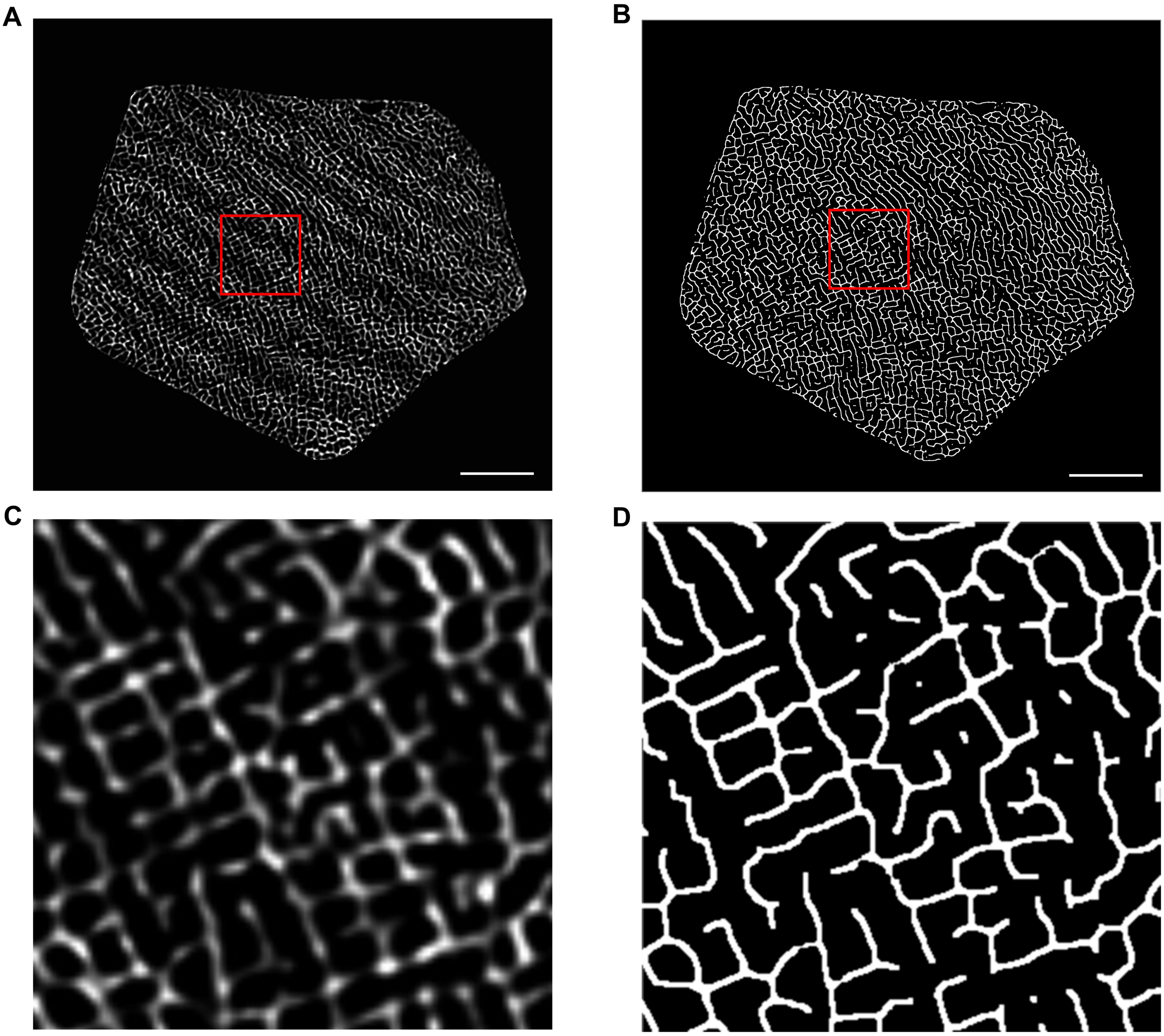


**Supplemental Figure 4. Using CellProfiler to determine the area occupied by the intermyofibrillar components.** (**A**) A representative image of an isolated muscle fiber before processing in CellProfiler. (**B**) Binary version of the image from A that was created by the custom “Intermyofibrillar Area” CellProfiler pipeline. The area occupied by the intermyofibrillar components is calculated from the area that is occupied by the white pixels in the binary image. (**C**-**D**) Zoomed-in images of the red-boxed regions in A and B. Scale bar = 10 μm

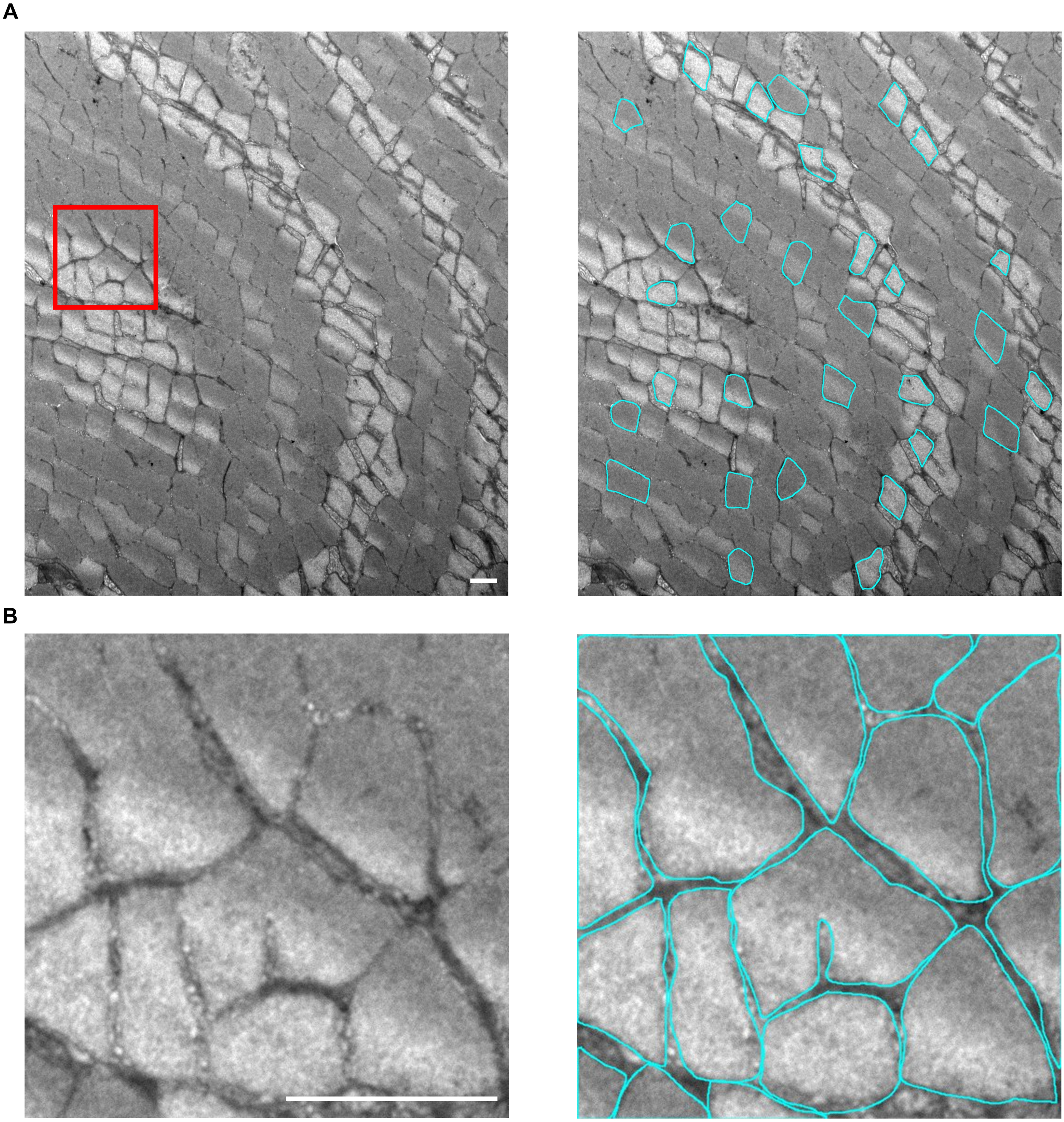


**Supplemental Figure 5. Manual tracing of myofibrils from images acquired using electron microscopy.** (**A**) Representative images show the manual tracing (cyan) of 30 randomly selected myofibrils for measurements of the average myofibril CSA. (**B**) Myofibrils within the red-boxed region of interest (ROI) were manually traced to obtain the CSA occupied by myofibrils within the ROI. The percentage of the fiber CSA occupied by myofibrils was calculated by dividing the CSA occupied by myofibrils within the ROI by the total area of the ROI and then multiplying by the fiber CSA. The number of myofibrils for the fiber was calculated by dividing the percentage of the fiber CSA occupied by myofibrils by the average myofibril CSA for that fiber (Scale bar = 1 μm).
