## Supplementary material for "A Novel Imaging Method (FIM-ID) Reveals that Myofibrillogenesis Plays a Major Role in the Mechanically Induced Growth of Skeletal Muscle": CellProfiler Supplemental Files: CellProfiler Pipeline User Instructions.pdf

### **Supplemental Methods**

Software

CellProfiler version 4.2.5 was downloaded from https://cellprofiler.org/previous-releases and used on a 64-bit Windows 10 PC (Intel Core i7-9850H CPU @ 2.60 GHz, 64 GB RAM). Images were processed prior to CellProfiler analysis using ImageJ (Fiji), which can be downloaded from https://fiji.sc. The pipelines for the quantification of myofibril cross-sectional area (Myofibril CSA Analysis) and the area occupied by SERCA1/2 + Autofluorescence (Intermyofibrillar Area), as well as example images for running these pipelines, are provided within the “CellProfiler Supplementary Files” folder.

Image Processing in ImageJ

We refer the reader to the main manuscript for the details of image acquisition and processing for both the mouse and human samples. The CellProfiler analyses described here are run the same for either tissue. For the purpose of this walkthrough, images of a mouse plantaris muscle that had been subjected to immunolabeling for SERCA1 are provided in the “Example Images” folder. The “Ch0_6144x6144_Image” file is a full-scale merged image of the SERCA1 and autofluorescence signals that had been subjected to deconvolution (see manuscript for more details). For processing, the “Ch0_6144x6144_Image” is opened with ImageJ. The “Rectangle ROI” tool is used to place a box around the fiber to be analyzed and then the “Crop” function under the “Image” tab is selected. These steps will create a cropped version of the original image.

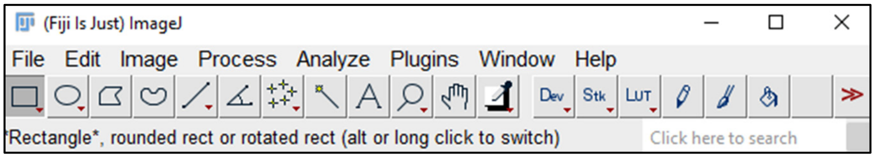


In the cropped version of the original image, the “Freehand Selection” tool is used to trace the periphery of the fiber of interest and then the fiber CSA is determined with the “Measure” function located under the “Analyze” tab (note: make sure that “Area” is selected under the “Set Measurements” function under the “Analyze” tab).

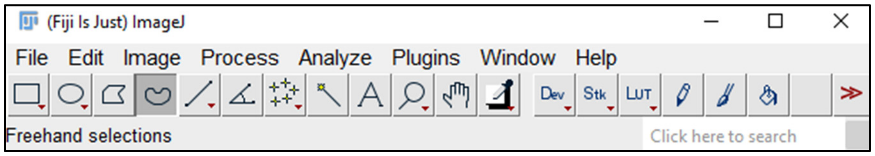


The fiber of interest is then isolated by using the “Clear Outside” function located under the “Edit” tab and saved as a .tiff file, making sure that “Ch0_” precedes the file name. For the purposes of this walkthrough, we have provided a file with an isolated fiber named “Ch0_Isolated_Fiber”, as well as a region of interest (ROI) from the fiber (“Ch0_ROI_of_Isolated_Fiber”). In this walkthrough, the image of the ROI (yellow box) will be used to demonstrate the functionality of the different CellProfiler pipelines.

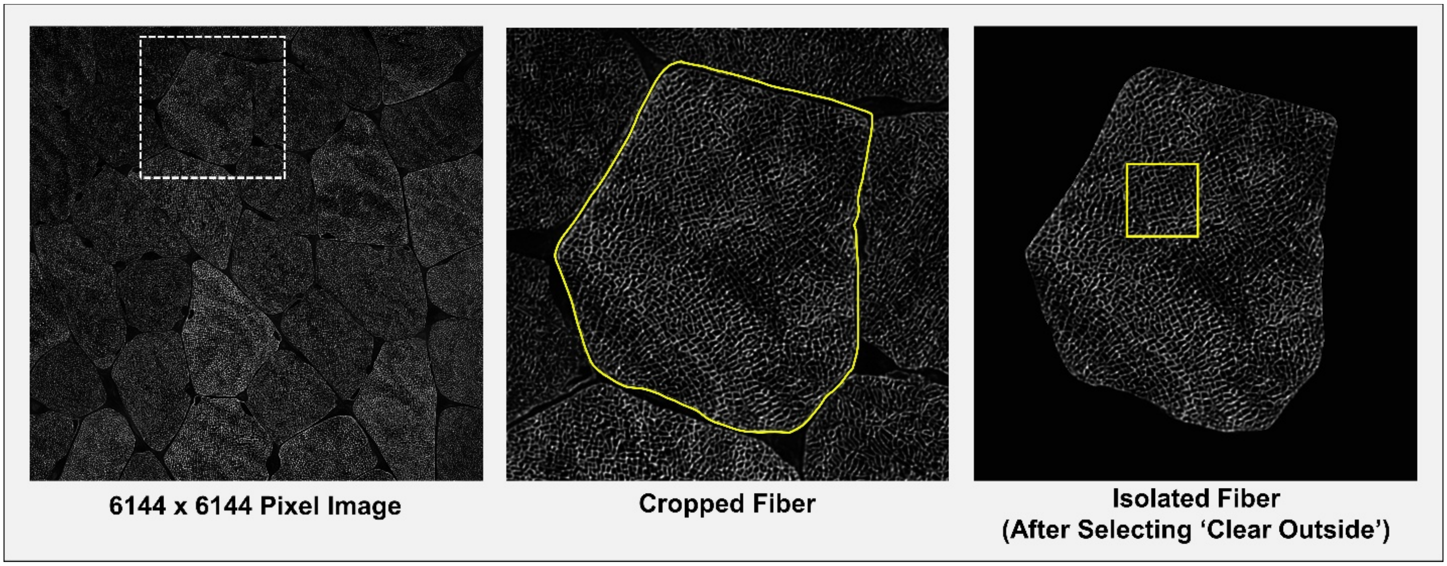


CellProfiler Pipeline and Image Loading

With CellProfiler version 4.2.5 installed on the computer, the pipeline of interest (e.g., "Myofibril CSA Analysis") can be opened by double-clicking on the pipeline in the "CellProfiler Supplementary Files" folder. The image(s) to be analyzed are then loaded by dragging the image(s) into the center box. Importantly, the program will recognize the images that are to be analyzed by looking for the "Ch0_" classifier. Any images that lack "Ch0_" will be excluded from the analysis.

The "Myofibril CSA Analysis" Pipeline

This pipeline is used to obtain measurements of myofibril size (i.e., CSA). After loading the image(s) to be analyzed, the program processes each image through 21 modules and exports the measurements of the qualified myofibrils as .csv files. The major functions of these different modules are described below.

*Image Processing*

In order to accurately identify individual objects (i.e., myofibrils), the images are first processed through 9 modules (RescaleIntensity, Closing, EnhanceOrSuppressFeatures, Closing, Threshold, Morph, ImageMath, Morph, Opening). These modules enhance and close minor gaps within the fluorescent signal. For illustrative purposes, the image "Ch0_ROI_of_Isolated_Fiber" was loaded into the pipeline, "Analyze Images" was clicked, and then the impact that each module had on the image is summarized below.

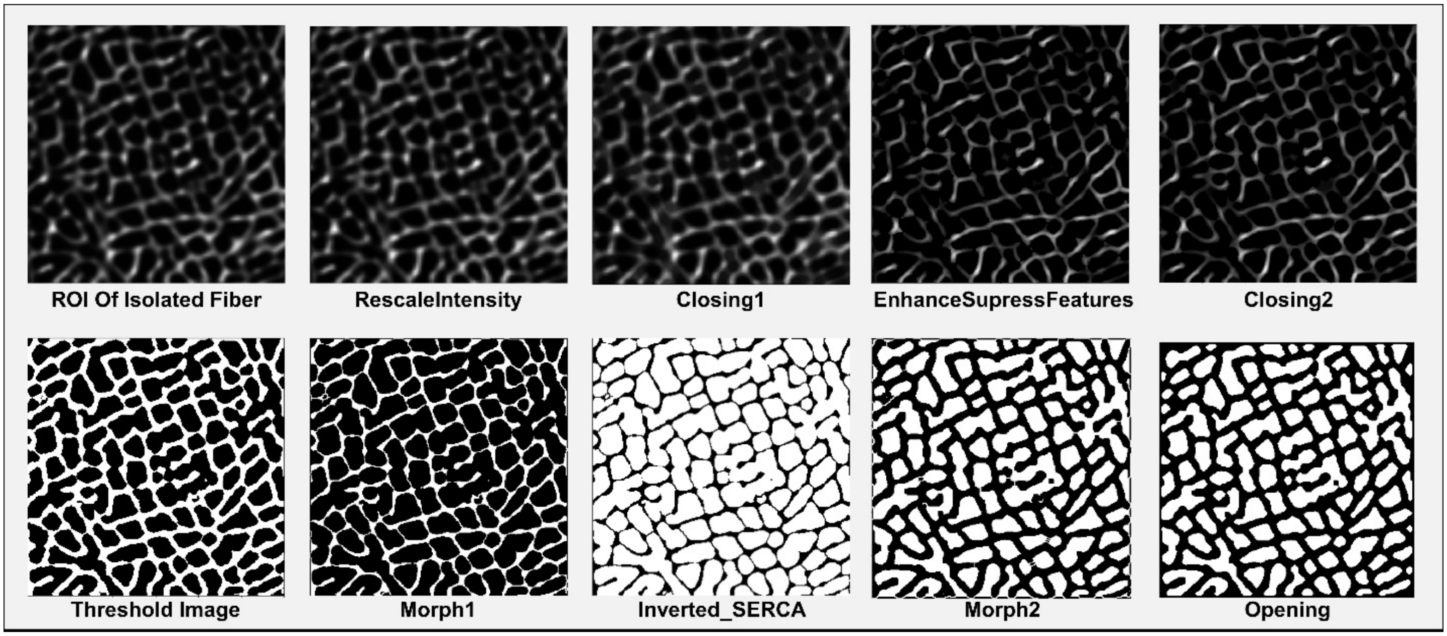

If users wish to see the impact of each of these modules in action, simply click the (eye) icon in the pipeline, and the impact of each selected module will be displayed during the image processing.

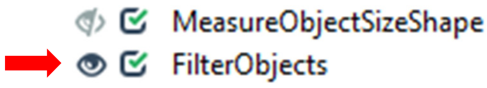


*Myofibril Identification and Filtering*

The next module "IdentifyPrimaryObjects" identifies objects based on the processed image. In this module, only objects that have diameters between 5-200 pixels are retained. Further filtering of objects is then performed in the "FilterObjects" module which removes all objects that do not meet the morphological criteria for subsequent measurements of myofibril size (see manuscript for details). Moreover, the cumulative impacts of the image processing etc. on the size of the myofibrils are corrected in the "ExpandOrShrinkObjects" module. Finally, an "EditObjectsManually" module is included so that, if desired, users can manually reject objects that should not be included in the final measurements. For instance, as shown on the next page, there are incomplete myofibrils that have been cropped off along the border of the ROI image, and these myofibrils can be removed prior to the measurements of myofibril size. To remove an object, the user clicks on the center of the outlined object, turning the solid outline of the object into a dashed line. The selection is then confirmed by clicking on "Done" on the bottom right (if no objects are to be removed, the user simply clicks "Done" for each image). After manually editing the filtered objects, measurements and images of the measured objects will be exported as described below.

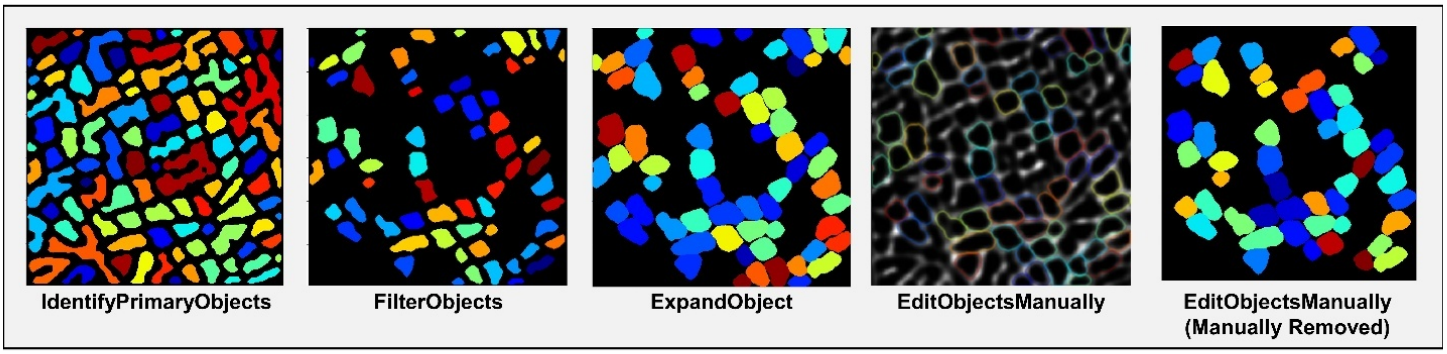


*Exported Results*

Measurements are determined with the "MeasureObjectSizeShape" module and exported via the "ExportToSpreadsheet" module. These modules produce separate .csv files with one containing measurements from each individual myofibril "Individual_Myofibril_CSA", while the other contains the average and standard deviation of the measurements for the entire image "Average_Myofibril_CSA". By default, the data will be saved to the users Desktop. If a different location is desired, the location can be selected within the "ExportToSpreadsheet" module under the "Output file location" section (shown below).

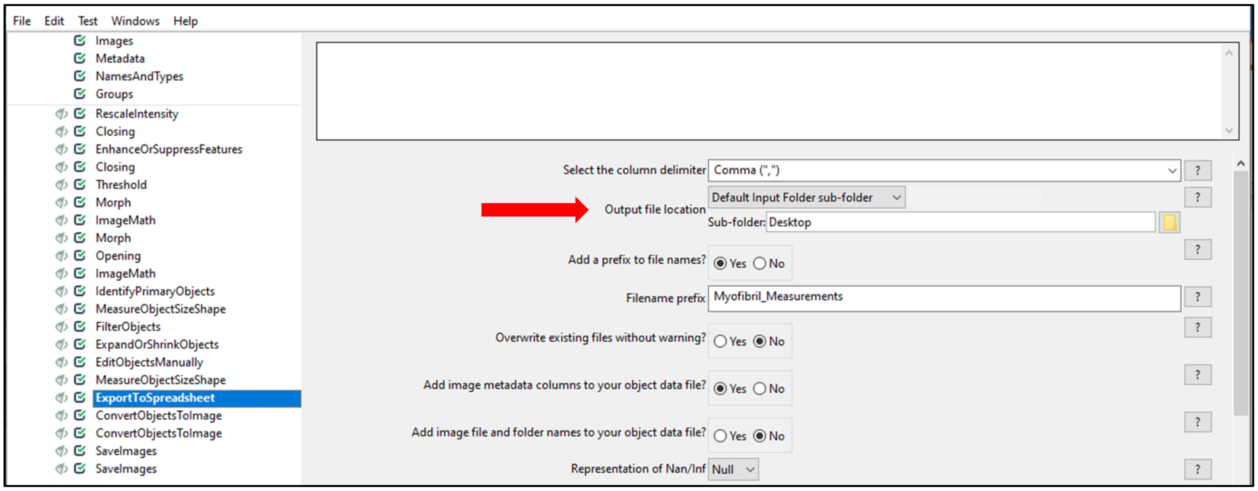


The final four modules convert the identified and filtered objects into images for visual evaluation of the filtering steps. By default, the images will be saved to the users Desktop. Similar to the process described above, if a different save location is desired, the location is chosen in both of the "SaveImages" modules under the "Output file location" section.

### The "Intermyofibrillar Area" Pipeline

This pipeline is used to obtain the measurements of the intermyofibrillar area. The image loading and processing steps are similar to the "Myofibril CSA Measurement" pipeline, but this pipeline measures the area of the signal from SERCA1/2 (and autofluorescence when applicable, see manuscript for details). Below are representative images of the processing steps of the "Ch0_ROI_of_Isolated_Fiber".

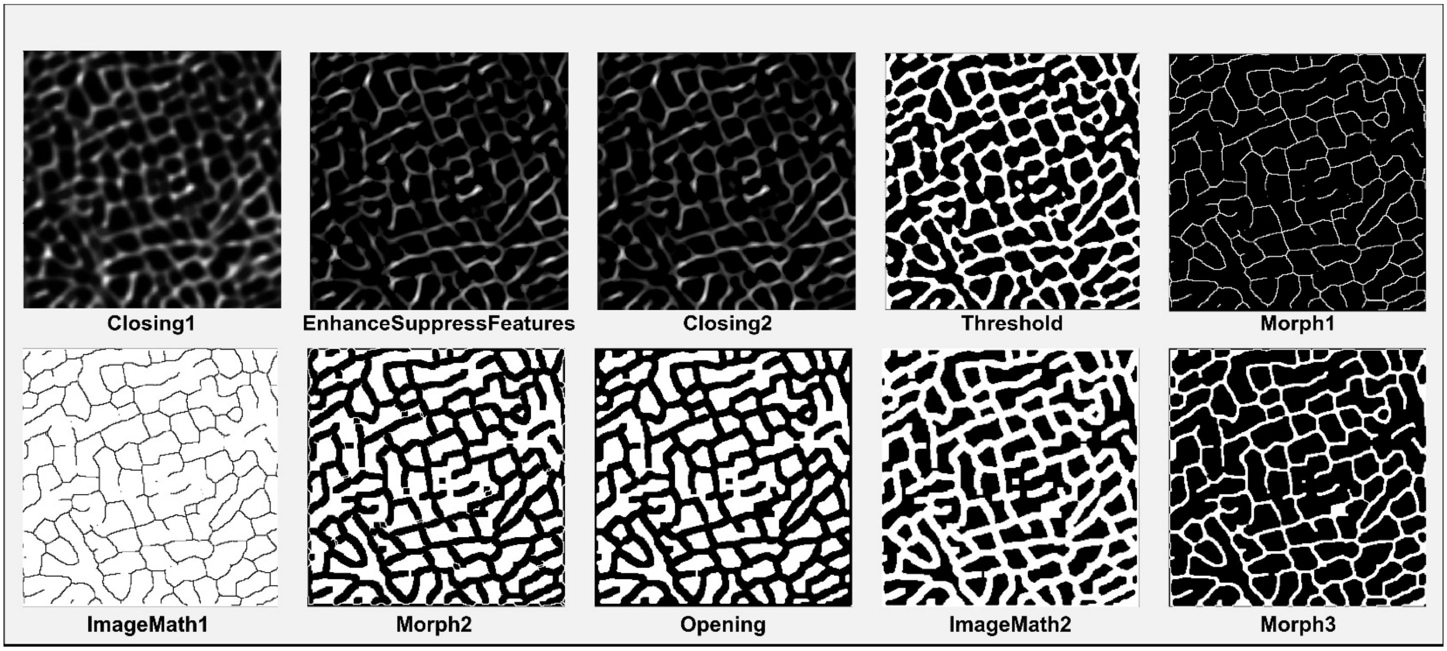


Measurements of the intermyofibrillar area are recorded with the module “MeasureImageAreaOccupied” and exported with the “ExportToSpreadsheet” module. By default, a single .csv file “Intermyofibrillar_Area” containing the intermyofibrillar area measurement for each image as well as a .tiff image of the identified intermyofibrillar area will be exported to the desktop. If a different save location is desired, the location is chosen in the “ExportToSpreadsheet” and “SaveImages” modules under the “Output file location” section. As illustrated below, a very high degree of overlap is observed when the image of the identified intermyofibrillar area is merged with the original signal.

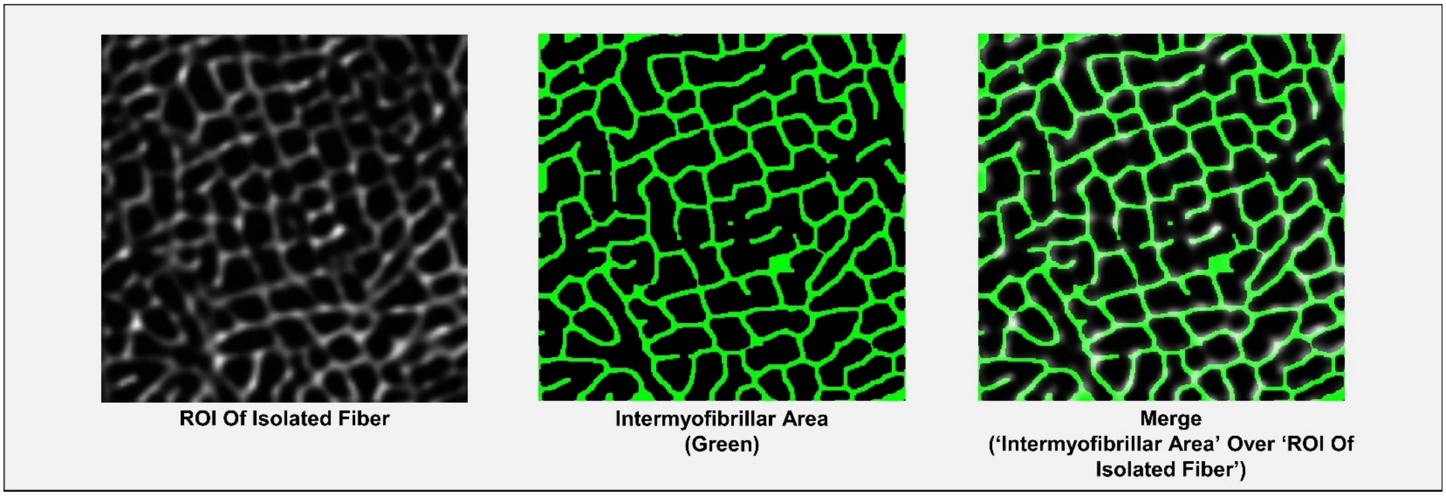


For illustrative purposes, we have demonstrated how the different modules perform on a small ROI within a single fiber. However, the “Myofibril CSA Analysis” and “Intermyofibrillar Area” pipelines are intended for use on whole muscle fibers. Illustrations of what the major outputs look like when a whole muscle fiber is analyzed are shown on the next page. These images were derived from the analysis of the “Ch0_Isolated_Fiber” image provided in the “Example Images” folder and can be reproduced by users to confirm that the pipelines are working appropriately. Additionally, other muscle fibers from the “Ch0_6144x6144_Image” can be selected and taken through the entire workflow to ensure that all steps are working appropriately.

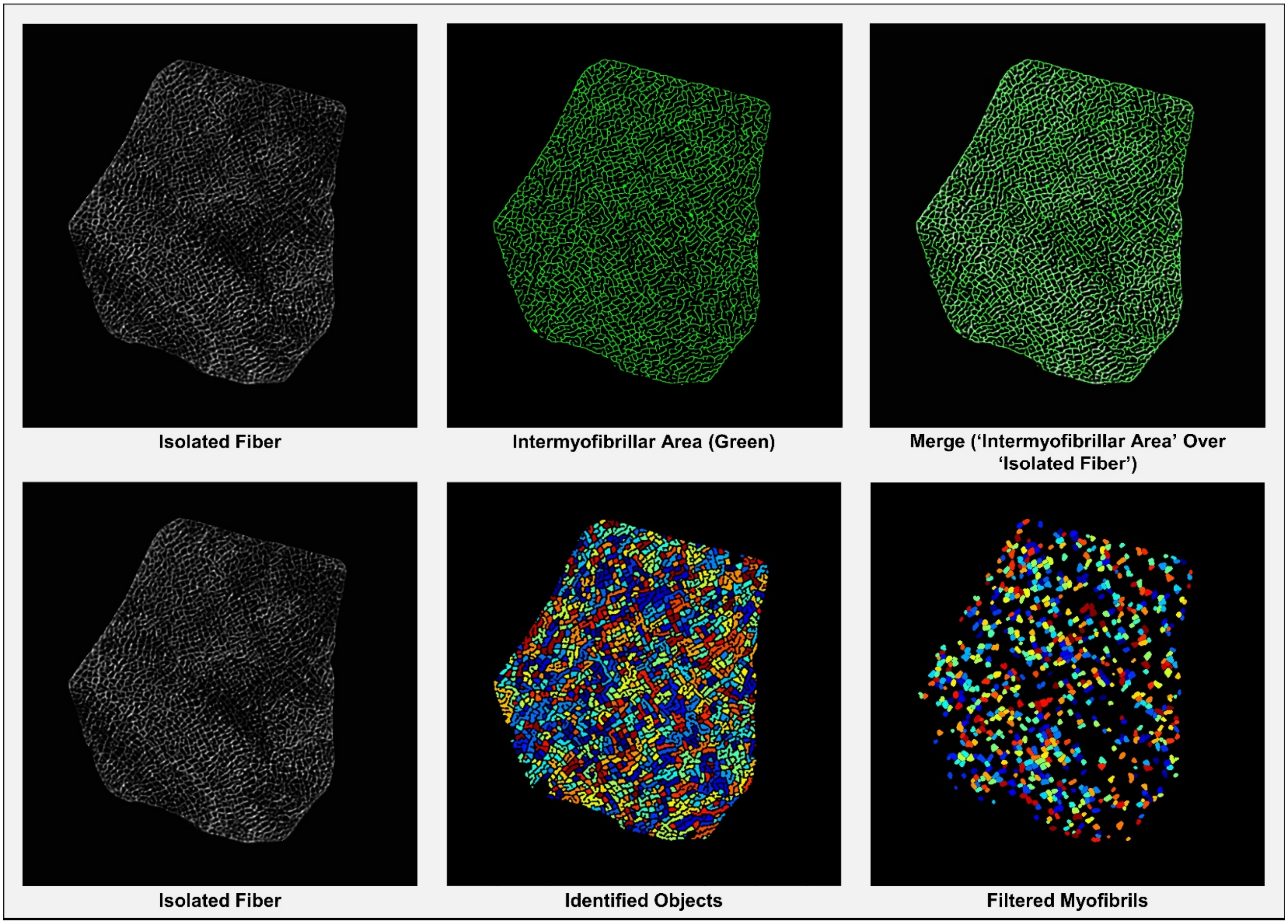


As a final note, the area measurements produced by the pipelines are exported in terms of the number of pixels$^2$. For the example images provided, a conversion of 844 pixel$^2$ per µm$^2$ should be used to convert the output to µm$^2$. Users who intend to analyze their own images will need to determine the appropriate conversion factor for their specific imaging conditions. Moreover, for the analysis of images that have been captured/processed with a system that is different from the one employed in this study, it is likely that minor modifications to the pipelines will be needed for optimal performance. Importantly, all settings in the pipelines can be modified and optimization will typically require an iterative trial-and-error process. Before modifying any settings, we strongly encourage users to familiarize themselves with the CellProfiler user manual which can be downloaded at https://cellprofiler-manual.s3.amazonaws.com/CellProfiler-4.2.5/index.html.
