## Supplementary figures and images for "A Novel Imaging Method (FIM-ID) Reveals that Myofibrillogenesis Plays a Major Role in the Mechanically Induced Growth of Skeletal Muscle"

### Ch0_Isolated_Fiber.tif

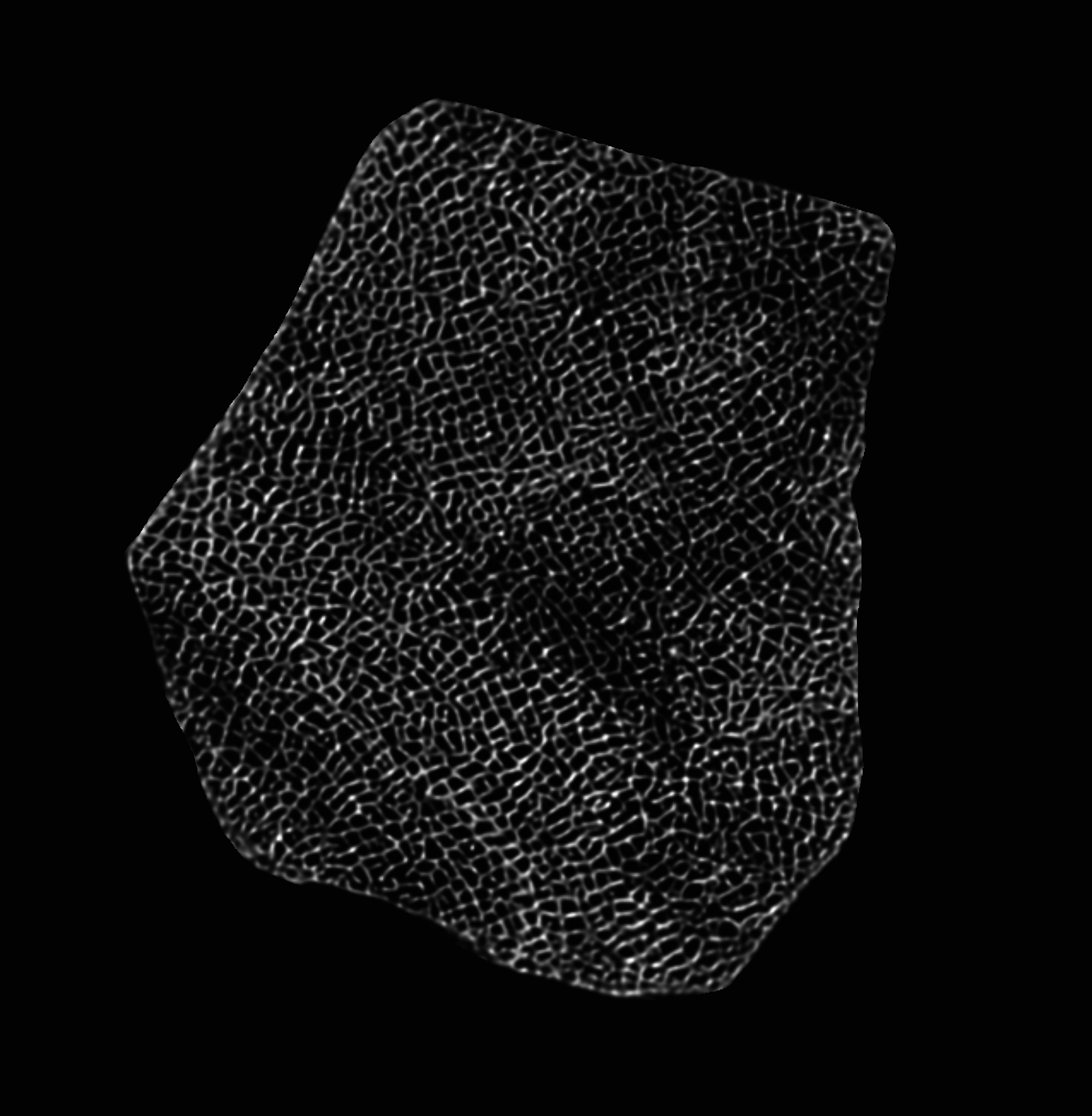

### Ch0_ROI_of_Isolated_Fiber.tif

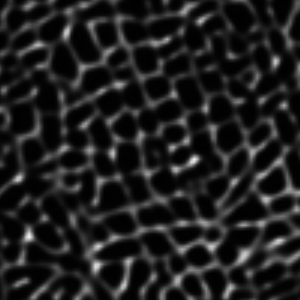
